# Overlapping and non-overlapping roles of the class-I histone deacetylase-1 corepressors, LET-418, SIN-3, and SPR-1 in *Caenorhabditis elegans* embryonic development

**DOI:** 10.1101/2020.07.21.213561

**Authors:** Yukihiro Kubota, Yuto Ohnishi, Tasuku Hamasaki, Gen Yasui, Natsumi Ota, Hiromu Kitagawa, Arashi Esaki, Muhamad Fahmi, Masahiro Ito

## Abstract

Histone deacetylases (HDACs) are divided into four classes. Class-I HDAC, HDAC-1 forms three types of complexes, namely the Nucleosome Remodeling Deacetylase complex, the Sin3 complex, and the CoREST complex, with specific corepressor component Mi2/CHD-3, Sin3, and RCOR1 in human, respectively. The functions of these HDAC-1 complexes are regulated by their corepressors, however, their exact mechanistic roles in several biological processes remain unexplored, such as in embryonic development. Here, we report that each of the corepressors, LET-418, SIN-3, and SPR-1, the homologous of Mi2, Sin3, and RCOR1, respectively, were expressed throughout *Caenorhabditis elegans* embryonic development and served essential roles in the process. Moreover, genetic analysis suggested that three pathways (i.e., LET-418– SIN-3–SPR-1, SIN-3–SPR-1, and LET-418) participated in embryonic development. Our terminal-phenotype observations of single mutants of each corepressor implied that LET-418, SIN-3, and SPR-1 played similar roles in promoting advancement to the middle and late embryonic stages. Genome-wide comparative-transcriptome analysis indicated that 47.5% and 42.3% of genes were commonly increased and decreased in *sin-3* and *spr-1* mutants, respectively. These results suggest that among the three pathways studied, the SIN-3–SPR-1 pathway mainly serves to regulate embryonic development. Comparative-Gene Ontology analysis indicated that these three pathways played overlapping and distinct roles in regulating *C. elegans* embryonic development.

## Introduction

During embryonic development, daughter cells generated from fertilized eggs contain the same genomic information as the mother cells when the cell-division process is completed. Although they have identical genome sequences, daughter cells can differentiate from precursor cells within developing tissues and organs through the epigenomic control of gene expression. Therefore, epigenomic modifications play important roles in normal embryonic development [1]. Epigenomic modifications are modulated via chemical changes to histones and DNA. Chemical acetylation and methylation of histones affect the regulation of gene expression by influencing histone– DNA and histone–protein interactions. Histone modifications are regulated by transferases and hydrolases [1]. Histone acetylation is positively regulated by histone acetyltransferase. Histone acetyltransferase promotes histone acetylation to neutralize the positive charge of the histone tail and, thus, acts as positive transcriptional regulator by weakening the physical interaction between the histone tail and DNA [2]. In contrast, histone deacetylase (HDAC) removes acetyl groups from histones and negatively regulates transcription by enhancing the physical interaction between the histone tail and DNA [3].

The 18 human HDAC proteins are divided into four classes based on their sequence homologies with the four yeast HDAC proteins [4, 5]. Class-I HDAC, HDAC-1 forms complexes with multiple components, such as transcriptional corepressors and DNA binding proteins, and promote histone deacetylation to suppress the transcription of target genes [6]. HDAC-1 transcriptional-corepressor can form three types of complexes in human, including the NuRD complex, the Sin3 complex, and the CoREST complex, where each complex includes a specific corepressor component (Mi2/CHD3, Sin3, and RCOR1, respectively) [6]. These complexes are thought to function as transcriptional repressors by inhibiting transcription of their target genes [6]. HDACs have been implicated in regulating various vital processes, such as DNA repair, lipid metabolism, cell-cycle progression, and the circadian rhythm [7-11]. Furthermore, HDAC-1 proteins have been shown to play important roles in the embryogenesis of multiple model organisms [12-15]. However, the mechanism whereby the three HDAC-1 complexes, NuRD, Sin3, and CoREST, participate in regulating embryonic development remains unknown.

The nematode *Caenorhabditis elegans* is a model multicellular organism, whose whole genome sequence and entire cell lineage have been completely identified [16, 17]. Therefore, *C. elegans* is a reliable model organism for analyzing the regulatory mechanism of embryogenesis. The constituents of the HDAC complex are also conserved in *C. elegans* [18]. The components of human HDAC-1 share conserved sequences with those of HDA-1 in *C. elegans*. The *C. elegans hda-1* gene can help regulate vulval development [19]. LET-418, SIN-3, and SPR-1 are *C. elegans* homologs of the human transcriptional corepressor components, Mi-2/CHD3, SIN3, and RCOR1, respectively, and each corepressor has been shown to be drive specific functions such as vulval development, male sensory cell formation, and gonadal morphogenesis during postembryonic development [20-23]. However, the functional relationships of these transcriptional corepressors in *C. elegans* embryogenesis remain unexplored.

In this study, we identified functional similarities and differences among the transcriptional corepressors, LET-418, SIN-3, and SPR-1, to understand the functional relationships of the three types of HDAC-1 corepressor in embryogenesis. First, we examined whether *hda-1, let-418, sin-3*, and *spr-1* participated in embryogenesis. Then, we analyzed the genetic interactions between two corepressors to identify relationships between all three corepressors. Finally, comprehensive comparative analysis of the target genes of the LET-418, SIN-3, and SPR-1 complexes was performed via RNA sequencing (RNA-Seq). We combined our analysis of genetic interactions with Gene Ontology (GO) analysis of these corepressors and found that the LET-418–SIN-3–SPR-1, SIN-3–SPR-1, and LET-418 pathways influenced embryonic development by positively and negatively regulating the associated pathway-specific gene functions. We also identified commonly regulated gene functions between the LET-418–SIN-3–SPR-1 and SIN-3–SPR-1 pathways.

## Materials and Methods

### *C*. *elegans* strains

*C. elegans* strains were derived from the wild-type (WT) Bristol strain [24]. Worms were incubated on nematode growth medium (NGM) and fed OP50 bacteria at 20°C. When performing RNA-interference (RNAi) experiments, the animals were fed RNAi bacteria, which were maintained at 20°C.

*C. elegans* strains with the following putative null alleles: *sin-3(tm1276*) (National BioResource Project, Japan), *spr-1(ok2144)* (*C. elegans* Gene Knockout Consortium), and the weak loss-of-function allele,*let-418 (n3536)* (Caenorhabditis Genetics Center) were used for our analysis.

### Sample preparation for RNA-Seq

To isolate synchronized early *C. elegans* embryos, the following four steps were performed. (1) Adult worms (WT and mutant) bearing fertilized eggs were treated with a bleach solution and the eggs were extracted. (2) The eggs were cultured in S-basal until all eggs hatched to synchronize the developmental stage, and subsequently OP50 solution was added to the S-basal. (3) The synchronized worms were incubated until they grew to the young-adult stage when they were capable of bearing 2–3 fertilized eggs. (4) The early embryos were isolated by bleaching.

### Total RNA extraction

For RNA-Seq and reverse-transcriptase quantitative polymerase chain reaction (RT-qPCR) analyses, total RNA was extracted from the WT, *let-418(n3536), sin-3(tm1476)*, and *spr-1(ok2144)* strains using the TRI Regent (Molecular Research Center, Inc., Cincinnati, OH). Following DNA digestion, the total RNA was extracted using an RNeasy Mini Kit (Qiagen). The extracted RNA was qualitatively evaluated using a Bioanalyzer (Agilent Technologies, Palo Alto, CA) and the Agilent RNA 6000 Nano Kit (Agilent Technologies, Palo Alto, CA).

### RT-qPCR analysis

Complementary DNA (cDNA) was synthesized from total RNA from WT *C. elegans* at each developmental stage (early-stage embryo, middle-stage embryo, late-stage embryo, first larva, and young adult) using the PrimeScript RT Reagent Kit (Takara). RT-qPCR was performed in a StepOnePlus(tm) qPCR system (Thermo Fisher Scientific) using THUNDERBIRD SYBR qPCR Mix (Toyobo). The expression levels of the *hda-1, sin-3, let-418*, and *spr-1* genes were normalized to that of a reference gene (Y45F10D.4), which was previously characterized as a reliable reference gene [25]. The following primers were used to amplify Y45F10D.1 (Y45F10D.4_F, 5′-GTCGCTTCAAATCAGTTCAGC-3′; Y45F10D.4_R, 5′-GTTCTTGTCAAGTGATCCGACA-3′), *hda-1* (hda-1_F, 5′-GGTCAAGGGCACGTCATGAAGCC-3′; hda-1_R, 5′-CTCGTCGCTGTGAAAACGAGTC-3′), *let-418* (let-418_F, 5′-GTGCTGCTATCGGATTGACAGACG-3′; let-418_R, 5′-GGGTTTGCCTCCAGTATTTGTGGC-3′), *sin-3* (sin-3_F, 5′-GCAACCGTGGAATTGATGA-3′; sin-3_R, 5′-GTTGATTCGGTGTTGTTCGAC-3′), and *spr-1* (spr-1_F, 5′-CTCCATCTCCATATCCTGAAGC-3′; spr-1_R, 5′-GCACGGCATTCTGGACGATTCATCG-3′).

### Feeding RNAi

RNAi was performed using the feeding-RNAi method with freshly prepared RNAi-feeding plates, as described [26]. Full-length *had-1, let-418, sin-3*, and *spr-1* cDNAs were isolated from a *C. elegans* cDNA library and inserted into the feeding-RNAi vector, L4440 (Addgene, Cambridge, MA, USA). An L4440 vector lacking an insert was used as a negative control. After confirming that each inserted sequence was correct, the feeding vectors were individually transformed into *Escherichia coli* HT115 (DE3) bacterial cells, which were then seeded on NGM**-**agar plates containing Luria–Bertani medium and 50 μg/mL ampicillin, and cultured for 12 h. Then, each culture was seeded onto a 60-mm feeding NGM agar plate containing 50 µg/mL ampicillin and 1 mM isopropyl β-D-1-thiogalactopyranoside, and then incubated at 25°C for 8 h. L4-stage worms were transferred onto a feeding plate and cultured at 20°C. The phenotypes of the F2 embryos were analyzed except for those fed RNAi bacteria that expressed double-stranded *hda-1* RNA, which were analyzed in the F1 embryos.

### Analysis of embryonic lethality

To analyze embryonic lethality, fertilized eggs were isolated by dissecting 1-day-old adult worms, after which the fertilized eggs were incubated at 20°C for 24 h and the ratio of the unhatched embryos were scored. To characterize the timing of the terminal phenotype, we defined embryonic lethality in early embryos as those that died before the ventral cleft-enclosure stage. Embryonic lethality in middle embryos was defined as those that died between the ventral cleft-enclosure stage and the comma stage. Embryonic lethality in late embryos was defined as those that died between the 1.5-fold stage and the 3-fold stage.

To analyze the terminal phenotypes of the dead embryos, Nomarski microscopy was performed using a Zeiss Axio Imager A1 microscope equipped with an EC Plan-Neofluar 40x NA, 0.75 objective (Zeiss), AxioVision software (Zeiss), and an AxioCam MRc digital camera. The images were processed using Adobe Photoshop CS6.

### RNA-Seq analysis

RNA-Seq analysis (N = 3) of the WT, *sin-3(tm1476), let-418(n3536)*, and *spr-1(ok2144)* strains was performed using a MiSeq instrument (Illumina), following the manufacturer’s recommended protocols (available on the Illumina website). Library preparation for RNA-Seq was performed using the TruSeq Stranded Total RNA LT Sample Prep Kit (Illumina). Next, the sample DNAs were denatured using a MiSeq Reagent Kit v3 (Illumina), diluted, and subjected to paired-end sequencing (75-base pair length) in a MiSeq instrument (Illumina).

### RNA-Seq data analysis

The quality of raw sequence data obtained by RNA-Seq was checked using FastQC software. Trimmomatic software [27] was employed to trim low-quality reads, and the sequence data were mapped to a *C. elegans* reference genome (WormBase Version 261) using HISAT2 software. The count data of wild type and mutants were compared with DESeq2 software [28] and differentially expressed genes (DEGs) (*p*-value < 0.01, log2 fold-change; positive or negative) were identified according to a previously described method [29]. To further analyze the DEGs, we identified up-regulated genes (log2 fold-change > 1 and *p*-value < 0.01) and down-regulated genes (log2 fold-change < −1 and *p*-value < 0.01). Using the DAVID Bioinformatics Resource database (version 6.8) [30]; GO enrichment analyses were performed to identify the specific functions of the DEGs.

### Statistical analyses of embryonic lethality

*P*-values (determined using Fisher’s exact test) were used to assess the significance of differences observed in terms of embryonic lethality. To analyze the embryonic lethality of *let-418(n3536);control(RNAi), let-418(n3536);sin-3(RNAi), let-418(n3536);spr-1(RNAi), sin-3(tm2376);control(RNAi), sin-3(tm2376);let-418(RNAi), sin-3(tm2376);spr-1(RNAi), spr-1(ok2114);control(RNAi), spr-1(ok2114);let-418(RNAi), spr-1(ok2114);sin-3(RNAi)*, the numbers of embryonic-lethal embryos and the number of hatched (non-embryonic-lethal) embryos were compared.

### Data availability

All data and samples described in this work will be freely provided upon request.

## Results

### Analysis of the mRNA-expression levels of *hda-1* and its corepressors, *sin-3, let-418*, and *spr-1* during development

*C. elegans* expresses two Mi2/CHD-3 homologs, LET-418 and CHD-3. Here, we only studied *let-418* to analyze the NuRD complex during the embryonic development of *C. elegans* because LET-418 is predominantly expressed during embryogenesis, in contrast to CHD-3 [31]. Changes in the relative mRNA-expression levels of *hda-1, let-418, sin-3*, and *spr-1* in *C. elegans* during development were analyzed by RT-qPCR at five developmental stages, including the early-embryo, middle-embryo, late-embryo, first-larval, and young-adult stages. We found that all of the analyzed genes were expressed beginning at the early-embryonic stage (Fig 1).

**Fig 1.**
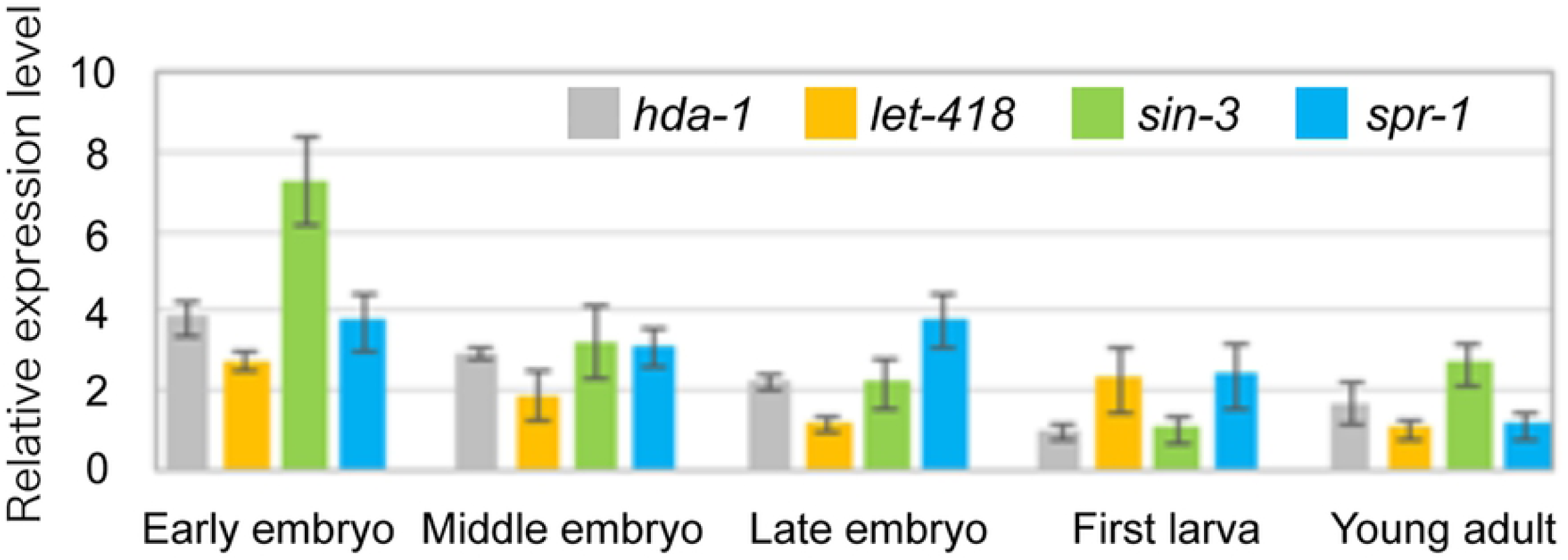
Comparison of the relative mRNA-expression levels of *hda-1, let-418, sin-3*, and *spr-1* during development. The mRNA-expression levels of *hda-1, let-418, sin-3*, and *spr-1* during the early-embryo, middle-embryo, late-embryo, first-larva, and young-adult stages in WT worms were analyzed by RT-qPCR (N = 3). The Y45F10D.4 gene was used as a reference. The bars and error bars indicate the relative mRNA-expression levels and standard deviations, respectively, of *hda-1* (gray), *let-418* (orange), *sin-3* (green), and *spr-1* (light blue) of mRNA at each developmental stage.

### Embryonic lethality of *hda-1(RNAi)* and mutants of the *hda-1* corepressors, *let-418, sin-3*, and *spr-1*

During *C. elegans* embryogenesis, the effects of HDA-1 and its corepressors were analyzed by observing the embryonic lethality of *hda-1(RNAi)* and *control(RNAi)* animals, the WT strain, and deletion (putatively null) mutants of *sin-3* and *spr-1*, and the temperature-sensitive weak allele of *let-418*. Embryonic lethality of the *let-418(n3536), sin-3(tm1276)*, and *spr-1(ok2144)* mutants (10.6%, 10.4%, and 5.3%, respectively) was much higher than that of the WT strain (1.1%). The embryonic lethality of *hda-1(RNAi)* animals, 99.7% (N = 352, data not shown), was much higher than that of the *control(RNAi)* animals, 4.6% (Fig 2).

**Fig 2.**
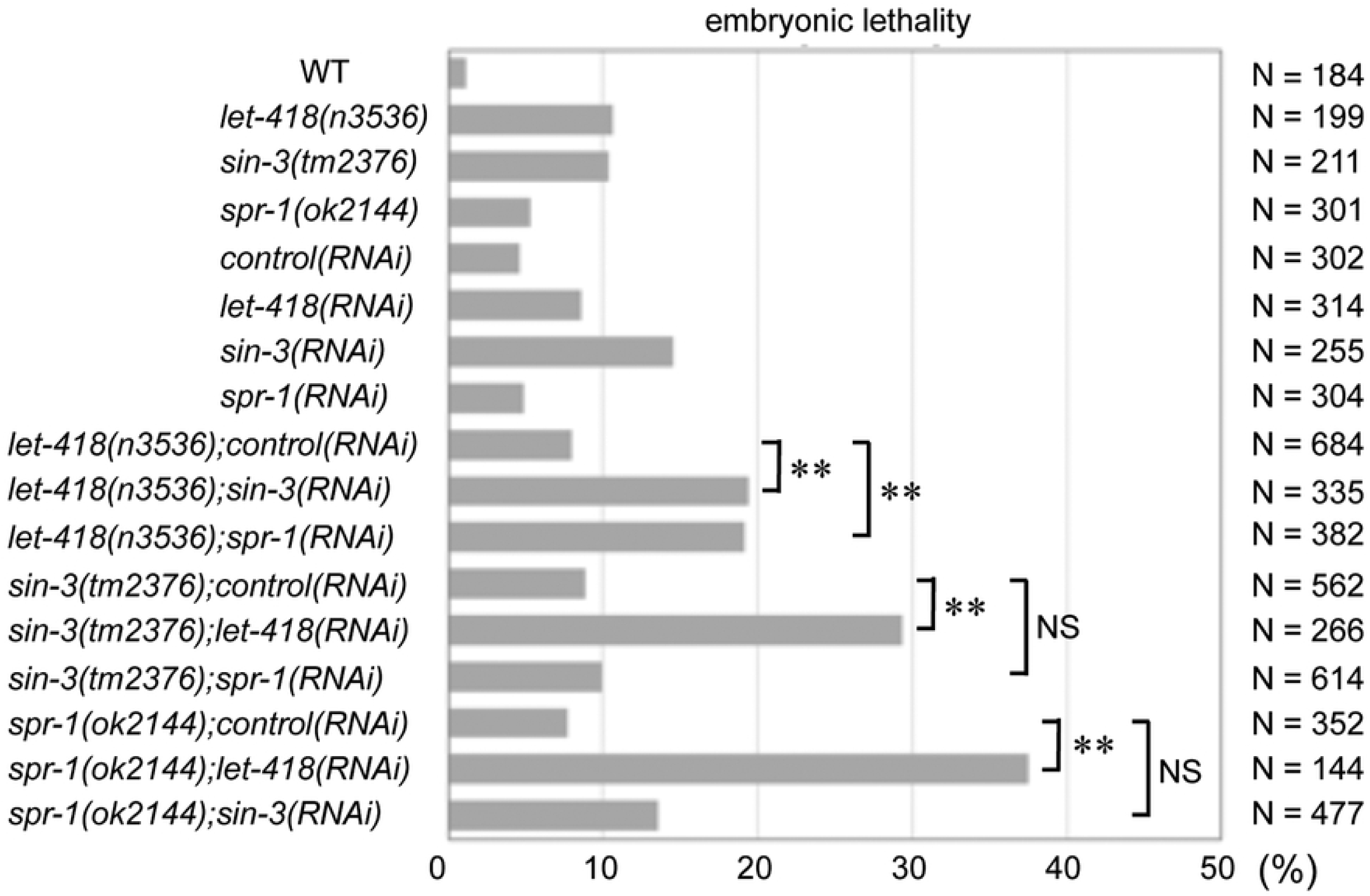
Genetic interaction of the *sin-3, let-418*, and *spr-1* mutants. Embryonic lethality of the WT strain and *let-418(n3536), sin-3(tm1476)*, and *spr-1(ok2144)* strains (single and double mutants) was analyzed by performing RNAi-based knockdown analysis. Each gray box indicates the embryonic lethality of the WT strain and indicated mutant. *P*-values are indicated for Fisher’s exact test comparison with *sin-3(tm1476); control (RNAi), let-418(n3536) control (RNAi)* or *spr-1(ok2144) control (RNAi)*. \*\**p*-value < 0.01. NS, not significant. N, number of embryos observed

### Genetic interactions among *let-418, sin-3*, and *spr-1* during embryonic development

To analyze functional relationships among *let-418, sin-3*, and *spr-1*, we conducted genetic analysis of the corepressors during embryonic development. To analyze genetic interactions among these corepressors, the effects of RNAi-mediated knockdown of one corepressor gene on other putative corepressor-null mutants were analyzed by calculating the resulting embryonic lethality. Interestingly, the embryonic lethality of the *sin-3(tm1276*);*spr-1(RNAi)* and *spr-1(ok2144);sin-3(RNAi)* mutants (9.9% and 13.6%, respectively) was almost comparable with those of *sin-3(tm1276);control(RNAi)* and *spr-1(ok2144);control(RNAi)* (8.9% and 7.7%, respectively). In contrast, the embryonic lethality of the *sin-3(tm1276*);*let-418(RNAi)* and *spr-1(ok2144);let-418(RNAi)* mutants (29.3% and 37.5%, respectively) was significantly higher than those of *sin-3(tm1276);control(RNAi)* and *spr-1(ok2144);control(RNAi)* (8.9% and 7.7%, respectively), as shown in Fig 2. These results suggest that *sin-3* and *spr-1* acted in same pathway during embryonic development. In contrast, *let-418* had function that was not shared with *sin-3* and *spr-1*.

To analyze genetic interactions between the *let-418(n3536)* weak mutant and the other corepressors, the effects of RNAi-based knockdown of the corepressor genes in the *let-418(n3536)* weak mutant were observed in terms of embryonic lethality. Interestingly, the embryonic lethality of the *let-418(n3536);sin-3(RNAi)* and *let-418(n3536);spr-1(RNAi)* mutants (19.4% and 19.1%, respectively) was significantly higher than that of *let-418(n3536);control(RNAi)*, which was 8.0% (Fig 2). These results imply the following two possibilities. (1) *let-418* participates in the same pathway as *sin-3* and *spr-1*. (2) *let-418* serves a function that does not overlap with that of *sin-3* and *spr-1* (Fig 2).

Taken together, these genetic analyses suggested that three pathways (i.e., the LET-418– SIN-3–SPR-1 pathway, the SIN-3–SPR-1 pathway) and the LET-418 pathway, regulate *C. elegans* embryonic development.

### Phenotypic analysis of the effects of the *let-418, sin-3*, and *spr-1* mutants on embryogenesis

The terminal phenotypes were observed using a differential-interference contrast microscope. Similar to *hda-1(RNAi)* embryos that were described previously [14], the development of most embryonic-lethal embryos stopped between the ventral cleft-enclosure stage to 3-fold stage in the *let-418(n3536), sin-3(tm1276*), and *spr-1(ok2144)* mutants (Fig 3, Table 1). These results indicate that each corepressor was essential for progression of the middle- and late-embryonic development stages.

**Table 1.** Timing of the embryonic lethality in *let-418, sin-3*, and *spr-1* mutants.

|  | Before ventral cleft enclosure (Early embryo) | Ventral cleft enclosure to comma (Middle embryo) | 1.5-fold to 3-fold (Late embryo) | Total number of embryos |
| --- | --- | --- | --- | --- |
| *let-418(n3536)* | 1 (2.4%) | 25 (61.0%) | 15 (36.6%) | 41 |
| *sin-3(tm2376)* | 2 (4.0%) | 36 (72.0%) | 12 (24.0%) | 50 |
| *spr-1(ok2144)* | 0 (0%) | 23 (65.7%) | 12 (34.3%) | 35 |

**Fig 3.**
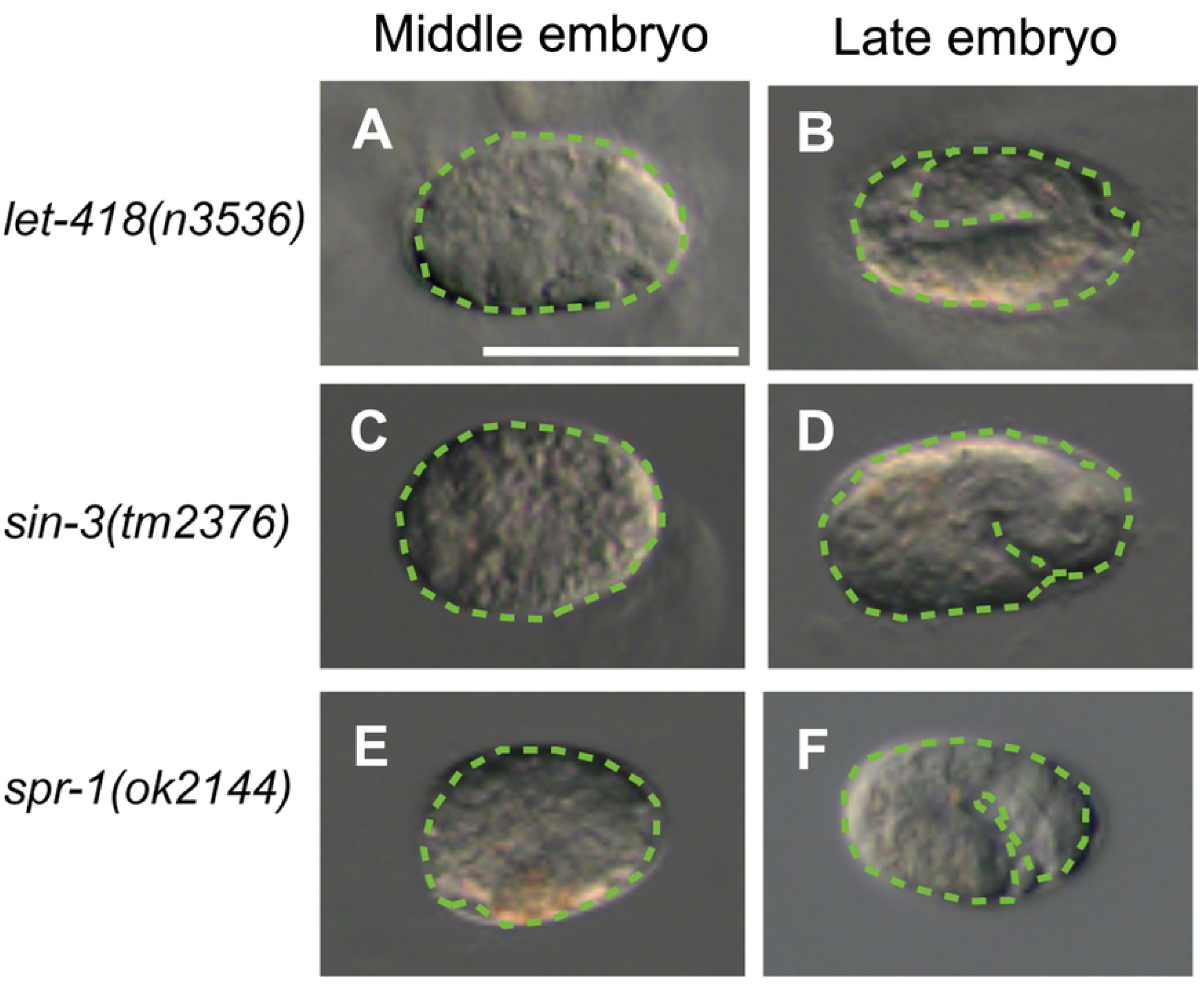
Microscopic images of the terminal phenotypes in *let-418, sin-3*, and *spr-1* mutants with embryonic lethality. Digital-interference contrast micrographs of the *let-418(n3536)* (A, B), *sin-3(tm1476)* (C, D), and *spr-1(ok2144)* (E, F) mutants. (A–F) The middle-stage (A, C, E) and late-stage (B, D, F) embryos that exhibited terminal phenotype of embryonic lethal are indicated. The green dotted lines outline the embryos in each panel. White scale bar, 50 μm.

### Identification of genes whose mRNA-expression levels were significantly different in the corepressor mutants

Expression-level information for 46,756 transcripts in early WT, *let-418(n3536), sin-3(tm1276)*, and *spr-1(ok2144)* mutants (N = 3) embryos was obtained by performing RNA-Seq analysis. DEGs in each mutant strain were defined as those whose expression levels significantly increased (*p*-value < 0.01 and log_2_-fold change > 1) or significantly decreased (*p*-value < 0.01 and log_2_-fold change < 1), when compared to the WT strain (Fig 4).

**Fig 4.**
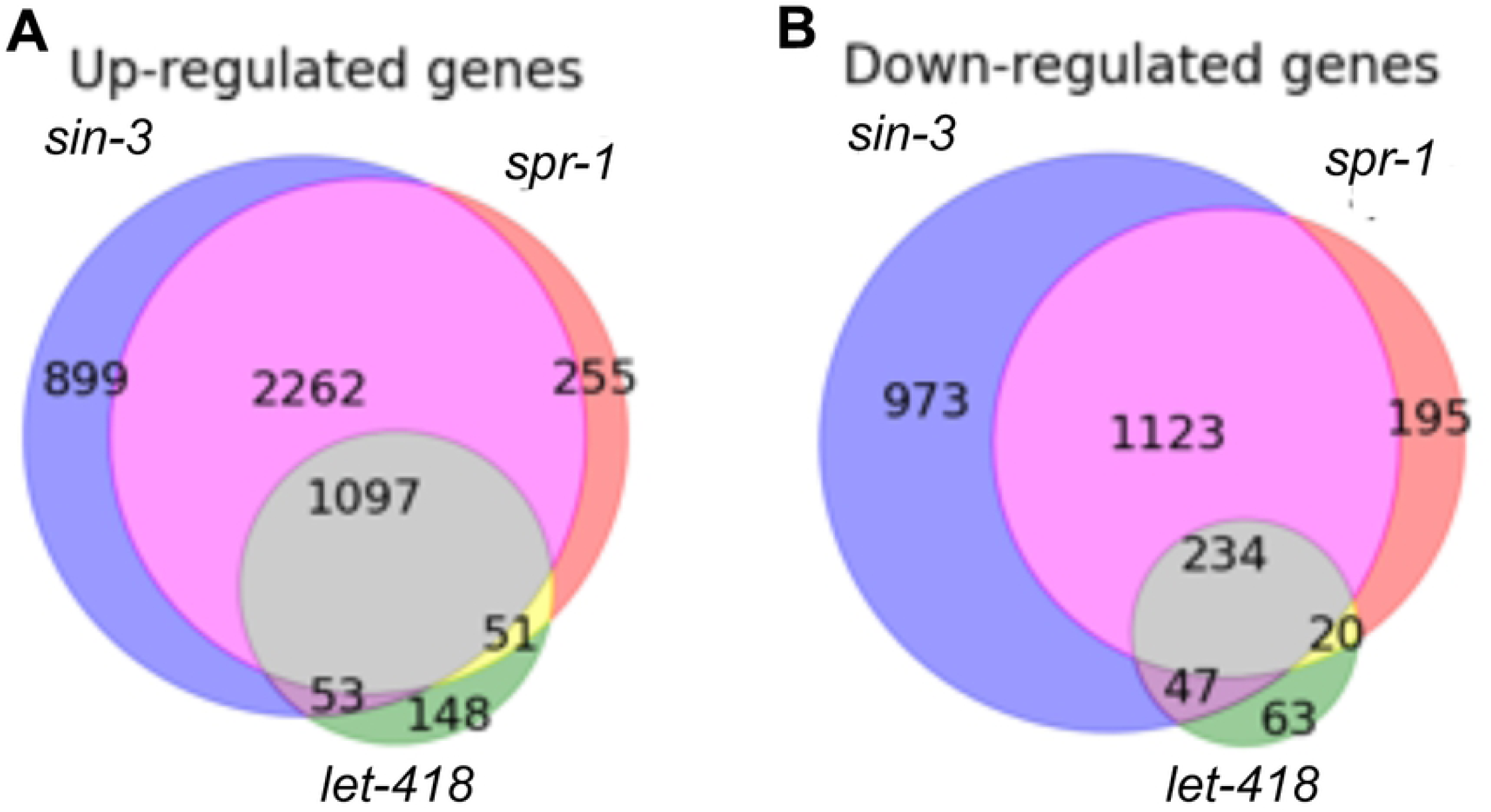
Comparison of mRNA-expression levels in corepressor mutants by RNA-Seq. Venn diagrams showing overlapping up-regulated genes (A) and down-regulated genes (B) in *let-418* embryos (green), *sin-3* embryos (blue), *spr-1* embryos (red), *let-418* and *sin-3* embryos (purple), *let-418* and *spr-1* embryos (yellow), *sin-3* and *spr-1* embryos (magenta) and *let-418, sin-3*, and *spr-1* embryos (gray).

### Analysis of transcriptionally regulated genes in *let-418, sin-3*, and *spr-1* mutants

To identify genes that were transcriptionally regulated by the three HDAC-1 complexes, we identified groups of genes whose expression levels significantly fluctuated in the corepressor mutants (Fig 4, S1–S3 Fig, S1 Table). Genes whose expression levels were significantly up-regulated (transcriptionally repressed) or down-regulated (transcriptionally promoted) in the corepressor mutants were defined as transcriptionally repressed and promoted genes, respectively (S2 Table). Based on these results, genes that were transcriptionally regulated by the three HDAC-1 complexes were classified into seven distinct, transcriptionally regulated gene groups (Fig 4). The frequencies of gene groups that were up-regulated by all three or more than one HDAC-1 corepressor mutants were 23.0% and 72.7%, respectively (Fig 4A). The frequencies of gene groups that were up-regulated in single mutants (*let-418, sin-3*, or *spr-1*) were 3.1%, 18.9%, and 5.4%, respectively (Fig 4A). The frequencies of gene groups that were up-regulated in double-corepressor mutants (*let-418* and *sin-3, let-418* and *spr-1*, or *sin-3* and *spr-1*) were 1.1%, 1.1%, and 47.5%, respectively (Fig 4A).

Similarly, the frequencies of gene groups that were down-regulated in all three mutants or in more than one HDAC-1 corepressor mutants were 8.8% and 53.5%, respectively (Fig 4B). The frequencies of gene groups that were down-regulated in single mutants (*let-418, sin-3*, or *spr-1*) were 2.4%, 36.6%, and 7.3%, respectively (Fig 4B). The frequencies of gene groups that were down-regulated in double-corepressor mutants (*let-418* and *sin-3, let-418* and *spr-1*, and *sin-3* and *spr-1*) were 1.8%, 0.8%, and 42.3%, respectively (Fig 4B).

We also checked which genes were significantly up-regulated and down-regulated in a single mutant (with mutation in *let-418, spr-1*, and *sin-3*; see S1–S3 Fig for the 10 most significantly up-regulated and down-regulated genes). Genes encoding extracellular matrix (ECM) components (*noah-1, lam-2* and *lam-3*), an ECM receptor (*dgn-1*), and a putative matrix proteinase inhibitor (*mig-6*) were among the most significantly up-regulated genes in the *let-418(n3536)* mutant. In the *sin-3(tm1276*) mutant, three ECM genes (*noah-1, lam-3*, and *nid-1*) as well as *mig-6* were significantly up-regulated. In addition, three ECM genes (*noah-2, lam-3*, and *nid-1*) and *mig-6* were up-regulated in the *spr-1(ok2144)* mutant. In contrast to the up-regulated genes, we did not identify any similarly down-regulated genes among the three mutants. These results indicated that all three class-I HDAC-1 corepressors significantly repressed the expression of ECM-related genes.

### GO enrichment analysis of transcriptionally regulated genes in the LET-418–SIN-3–SPR-1, SIN-3–SPR-1, and LET-418 pathways

HDAC-1 complexes are known to function as negative regulators of transcription. GO enrichment analysis was performed with the seven groups of up-regulated (transcriptionally repressed) genes (Fig 4A and S3 Table) to gain further insight into their potential roles in *C. elegans* embryogenesis.

To characterize the biological roles of the LET-418–SIN-3–SPR-1, SIN-3–SPR-1 pathway, and LET-418 pathways, we focused on differences and similarities in GO terms related to embryogenic development, cell specification, cell differentiation, cellular function, gene expression, and molecular function among the *let-418, sin-3*, and *spr-1* mutants, the *sin-3* and *spr-1* mutants, and the *let-418* mutant. First, we focused on similarities in GO terms associated with the up-regulated genes among the three groups. Various GO terms such as nervous system development, cell adhesion, epithelium/epithelial cell development and muscle cell development, and actin cytoskeleton/actomyosin structure organization were found between all three mutants and between the *sin-3* and *spr-1* mutants (Fig 5A, B). Next, we focused on GO terms that were specifically identified in these pathways. When we focused on genes specifically up-regulated in all three mutants, we identified several associated GO terms, including embryonic morphogenesis, cell fate commitment, cell migration, and positive regulation of gene expression (Fig 5A). Up-regulated genes in both the *sin-3* and *spr-1* mutants were associated with various GO terms, such as cilium morphogenesis, ion transport, cell morphogenesis, nervous system development, cell-cell signaling, establishment of localization along microtubule, and regulation of cell communication (Fig 5B). Although genes up-regulated in the *let-418* mutant were associated with GO terms such as lipid storage, transmembrane transport, and intracellular signal transduction, the *p*-values did not reflect significant differences (Fig 5C).

**Fig 5.**
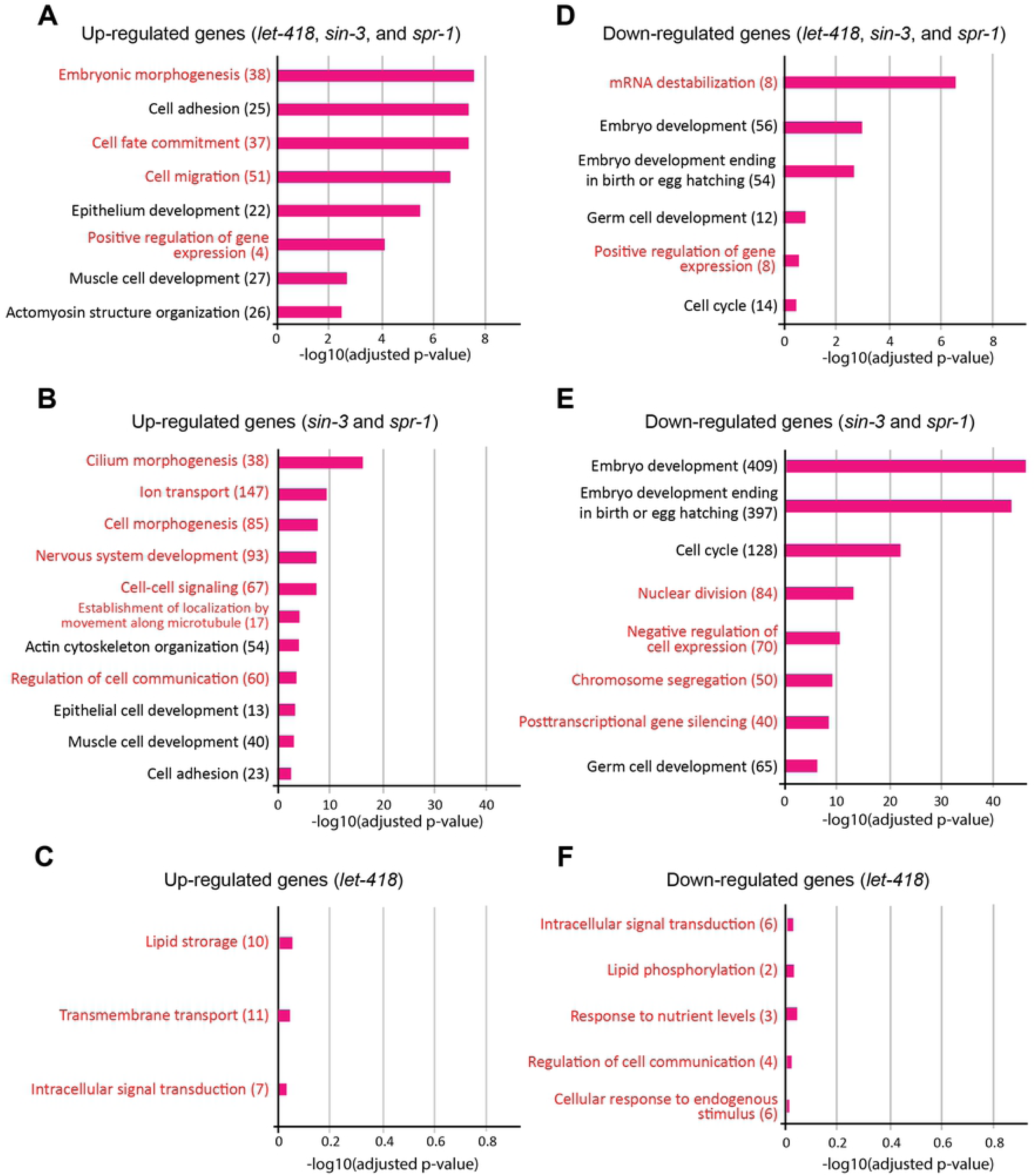
GO analysis of genes dysregulated by the SIN-3–SPR-1–LET-418, SIN-3– SPR-1, and LET-418 pathways. Go term of the gene groups that were elucidated by up-regulated genes (A-C) and down-regulated genes (D-F) from the corepressor mutants were indicated. (A) GO terms associated with genes that were transcriptionally repressed by the LET-418–SIN-3–SPR-1 pathway. (B) GO terms associated with genes that were transcriptionally repressed by the SIN-3–SPR-1 pathway. (C) GO terms associated with genes that were transcriptionally repressed by the LET-418 pathway. (D) GO terms associated with genes that were transcriptionally promoted by the LET-418–SIN-3–SPR-1 pathway. (E) GO terms associated with genes that were transcriptionally promoted by the SIN-3–SPR-1 pathway. (F) GO terms associated with genes that were transcriptionally promoted by the LET-418 pathway. The terms indicated with black text were identified as common GO terms between the LET-418–SIN-3–SPR-1 pathway and the SIN-3–SPR-1 pathway. The red text indicates specific GO terms associated with the LET-418–SIN-3–SPR-1 pathway, the SIN-3–SPR-1 pathway, and the LET-418 pathway. The p-values of the GO terms were determined to evaluate the potential relevance of the associated biological pathways. The numbers in each set of parentheses indicate the numbers of genes that were associated with each GO term.

When we focused on down-regulated (transcriptionally promoted) genes (Fig 4B), similar GO terms, embryo development, embryo development ending in birth or egg hatching, germ cell development, and cell cycle were enriched in association with the all three mutants and with both of the *sin-3* and *spr-1* mutants (Fig 5D, E). When we focused on genes that were specifically down-regulated in all three mutants, various GO terms such as mRNA destabilization and positive regulation of gene expression were found (Fig 5D). Genes specifically down-regulated in both the *sin-3* and *spr-1* single-mutants were associated with various GO terms such as nuclear division, negative regulation of gene expression, chromosome segregation, and posttranscriptional gene silencing (Fig 5E). Although genes specifically down-regulated in *let-418* mutant were associated with GO terms such as cellular response to endogenous stimulus, intracellular signal transduction, response to nutrient levels, lipid phosphorylation, and regulation of cell communication, the *p-*values did not indicate that these associations were statistically significant (Fig 5F).

## Discussion

In this study, we found that all three HDAC-1 corepressor gene mutants (*sin-3, let-418*, and *spr-1*) exhibited embryonic lethality. Similar to previous phenotypic observations with *hda-1(RNAi)-*knockdown embryos [14], the *let-418, sin-3*, and *spr-1* single-mutants showed embryonic lethality between the middle-to late-embryonic stages. These results suggested that three types of HDAC-1 complexes, namely the NuRD, Sin3, and CoREST complexes serve important roles in *C. elegans* embryogenesis. Consistent with previous results obtained with *C. elegans*, zebrafish, and mice [32-34], we found that *hda-1* was expressed during the early-embryonic stage. Moreover, the HDAC-1 corepressors, *let-418, sin-3*, and *spr-1* showed significantly elevated expression at the early-embryonic stages of the WT strain. These results imply that all three types of HDAC-1 complexes (i.e., the LET-418, SIN-3, and SPR-1 complexes) may regulate embryonic development beginning at the early-embryonic stage. Genetic analysis of these corepressors indicated that the LET-418–SIN-3–SPR-1, SIN-3–SPR-1, and LET-418 pathways were involved in regulating embryogenesis. However, the functional relationships between SIN-3 and SPR-1 in the SIN-3–SPR-1 pathway and those among LET-418, SIN-3, and SPR-1 in the LET-418–SIN-3–SPR-1 pathway remain uncharacterized. The terminal phenotypes of the *sin-3* and *spr-1* null mutants and the *let-418* weak mutant were similar to each other, and therefore it was difficult to determine the epistatic relationships among these corepressors. Further studies are required to identify the signal-transduction cascades activated by these corepressors in each pathway.

Regarding the LET-418 complex, it has been shown that *let-418* mRNA and the encoded protein are ubiquitously expressed in early-embryonic cells [31]. However, among the seven gene groups analyzed in this study less genes were specifically repressed by the LET-418 complex than by the other two complexes. One possible explanation for this result is that the LET-418 complex may be regulated by the other complexes. Another possibility is that a low number of target genes are specifically down-regulated by the LET-418 complex during early embryogenesis. Taken together, these results imply that although LET-418 significantly contributes to embryogenesis, LET-418 transcriptionally targets relatively few genes during embryogenesis. The *let-418* mutant used in this study is weak allele of the *let-418* gene, and therefore the transcriptome analysis does not fully reflect the normal function of this gene during embryonic development.

Next, a genome-wide comparative analysis of gene expression was performed by RNA-Seq. The percentage of genes transcriptionally repressed by more than two types of HDAC-1 complexes (i.e., SIN-3–SPR-1 and LET-418–SIN-3–SPR-1 pathways; 72.7%) was higher than the combined percentage of genes transcribed independently by the three HDAC-1 complexes (27.3%). These results imply that the three types of HDAC-1 complexes cooperatively controlled early embryonic development via histone deacetylation. For example, the SIN-3 and SPR-1 corepressors transcriptionally repressed a high percentage (47.5%) of the same target genes, and SIN-3 and SPR-1 play similar roles in regulating the expression of genes that regulate embryonic development. Similarly, all three corepressors were found to transcriptionally repress a moderate percentage (23.0%) of the same target genes. One possible explanation of these results is that both SIN-3 and SPR-1, as well as all three corepressors, serve overlapping roles in regulating gene expression. Therefore, we speculate that because HDAC-1 corepressors share the same target genes during embryogenesis, functional redundancy among the corepressors may occur during *C. elegans* embryogenesis.

Comparative GO analysis among the three pathways indicated that similar GO terms were enriched between the LET-418–SIN-3–SPR-1 pathway and the SIN-3–SPR-1 pathway, but not the LET-418 pathway. These results imply that cooperative regulation of gene expression among the three corepressors and between SIN-3 and SPR-1 are important for precise regulation of embryonic development. Further analyses of the GO terms indicated that many of the suppressed genes were related to neuronal, epithelial, and muscle development and actin-structure regulation. Enhanced expression of genes related to embryonic and germ cell development, and cell-cycle progression was identified as a common feature between the LET-418–SIN-3–SPR-1 and SIN-3–SPR-1 pathways (Fig 6).

**Fig 6.**
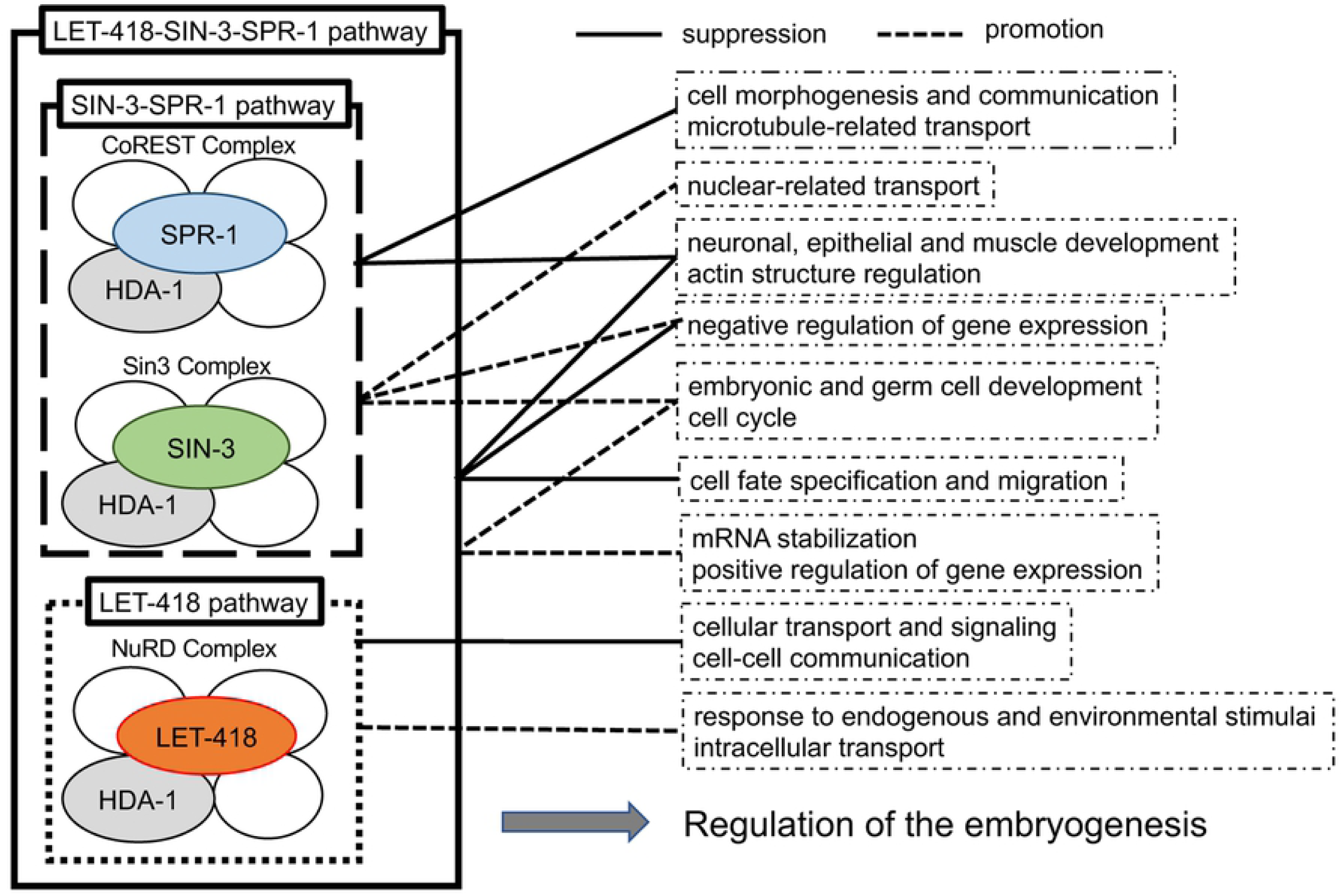
Model of the functional relationships among the LET-418–SIN-3–SPR-1, SIN-3–SPR-1, and LET-418 pathways. The LET-418–SIN-3–SPR-1, SIN-3–SPR-1, and LET-418 pathways positively and negatively regulate common and pathway-specific biological functions to influence embryonic development.

Similar to previous findings indicating that ECM genes were up-regulated in *hda-1(RNAi)* embryos [35], we found that genes encoding ECM and ECM-related mRNAs were significantly up-regulated in the HDAC-1 corepressor mutants, *let-418(n3536)*, sin-3*(tm1476)*, and *spr-1(ok2144)*. The up-regulated ECM genes (*noah-1, noah-2, nid-1, lam-2*, and *lam-3*) and ECM-related genes (*dgn-1* and *mig-6*) play important roles in embryonic morphogenesis or neuronal patterning [36-41], and therefore their temporal suppression is important for the regulation of embryonic development. Thus, we speculate that all three corepressors serve a common role that is required for the negative regulation of genes encoding ECM and ECM-related proteins.

While focusing on gene suppression specific to the SIN-3–SPR-1 pathway, we identified genes related to cell morphogenesis, intracellular communication, and microtubule-related transport that were specifically repressed (Fig 6). In contrast, when we focused on the promotion of gene expression, we identified genes related to nuclear-related cell division and negative regulation of gene expression that were promoted (Fig 6). Interestingly, mSin3A and CoREST were co-expressed in mouse embryos at E11.5, and mSin3A has been shown to act as a functional component of the REST–CoREST suppressor complex [42]. Thus, negative transcriptional regulation of the SIN-3–CoREST suppressor complex might be conserved both in vertebrates and invertebrates.

Suppressive effects that were specific to the LET-418–SIN-3–SPR-1 pathway identified repression of genes related to cell morphogenesis, cell-fate specification, cell migration, and negative regulation of gene expression (Fig 6). Although *hda-1* has been shown to positively regulate neuronal and distal-tip cell migration during post embryonic development in *C. elegans* [32, 43], negative regulation of cell migration by the LET-418–SIN-3–SPR-1 pathway was specifically regulated during embryonic development. In contrast, mRNA stabilization and positive regulation of gene expression were promoted. Differences in gene expression and mRNA stabilization occurred in somatic and germ cell linages throughout embryonic development in *C. elegans*, which may reflect the function of the LET-418–SIN-3–SPR-1 pathway. Taken together, our results indicate that LET-418–SIN-3–SPR-1 positively and negatively regulated cell-type-specific functions during embryogenesis. Thus, all three HDA-1 corepressor complex (the LET-418, SIN-3, and SPR-1 complexes) cooperated to negatively regulate cell differentiation and movement, promote cell-type-specific gene stabilization, and positively and negatively regulate the expression levels of different genes.

When focusing on suppressive effects specific to LET-418 pathway, we identified genes related to controlling cellular transport and signaling that were specifically repressed. In contrast, genes related to controlling cell–cell communication, cellular responses to endogenous and environmental stimuli, and intracellular transport were promoted (Fig 6). Here, we used a weak allele of the *let-418* mutant, and further studies with a strong loss-of-function *let-418* mutant are required to confirm its normal cellular roles.

## Conclusions

Using combined analyses of genetic interactions and transcriptome levels, we identified cooperative functions among the *C. elegans* homologs of the HDAC-1 corepressors, LET-418, SIN-3, and SPR-1. Genetic analyses suggest that three pathways (i.e., the SIN-3–SPR-1–LET-418, SIN-3–SPR-1, and LET-418 pathways) play important roles in embryonic development via transcriptional regulation. We also found that similar to the *hda-1(RNAi)* embryos that have been previously described, embryonic lethality occurred between the middle- and late-embryonic stages. Finally, comparative RNA-Seq analysis of these three pathways indicated that approximately half of up-regulated and down-regulated genes were associated with the SIN-3–SPR-1 pathway. Similarly, 10– 20% of the up-regulated and down-regulated genes were associated with the LET-418– SIN-3–SPR-1 pathway. Taken together, our findings suggest that the class-I HDAC-1 corepressors, LET-418, SIN-3, and SPR-1 cooperatively regulate embryogenesis by positively and negatively regulating gene-expression levels. We speculate that cooperative functions among HDAC-1 corepressors might be important for precise regulation of embryonic development both vertebrates and invertebrates.

## Acknowledgements

We would like to thank Mr. Yusuke Nomoto and Mr. Takahiro Nakamura for their support and helpful comments. Some strains were provided by the CGC, which is funded by NIH Office of Research Infrastructure Programs (grant number P40 OD010440), the *C. elegans* Gene Knockout Consortium, and the National Bioresource Project in Japan (lead by S. Mitani).

## Supporting Information

**S1 Fig. Volcano plot of the *let-418(n3536)* mutant versus the WT strain, highlighting the 10 most significantly up-regulated and down-regulated genes related to embryogenesis**. The blue and red dots indicate down-regulated and up-regulated genes, respectively. A *p*-value of < 0.05 was used as the threshold for statistical significance.

**S2 Fig. Volcano plot of the *sin-3(tm1276)* mutant versus the WT strain, highlighting the 10 most significantly up-regulated and down-regulated genes related to embryogenesis**. The blue and red dots indicate down-regulated and up-regulated genes, respectively. A *p-*value of < 0.05 was used as the threshold for statistical significance.

**S3 Fig. Volcano plot of the *spr-1(ok2144)* mutant versus the WT strain, highlighting the 10 most significantly up-regulated and down-regulated genes related to embryogenesis**. The blue and red dots indicate down-regulated and up-regulated genes, respectively. A *p*-value of < 0.05 was used as the threshold for statistical significance.

**S1 Table. List of the dysregulated genes observed in the *let-418(n3536), sin-3(tm1276)*, and *spr-1(ok2144)* mutants, compared to the WT strain**. Log2-normalized RNA-Seq data are shown, indicating the frequencies of dysregulated genes in the *let-418(n3536), sin-3(tm1276)*, and *spr-1(ok2144)* mutants, divided by the corresponding expression levels in the WT strain. A *p*-value of 0.01 was used as the cut-off.

**S2 Table. List of up-regulated and down-regulated genes in the *let-418, sin-3*, and *spr-1* mutants**. Expression levels (based on the RNA-Seq data) are shown for up-regulated genes (log2 fold-change > 1 and *p*-value < 0.01) and for down-regulated genes (log2 fold-change < −1 and *p*-value < 0.01).

**S3 Table. GO terms that were associated with the three pathways studied**. GO terms associated with up-regulated and down-regulated genes that participate in embryogenic development, cell specification, cell differentiation and cellular function, gene expression, and molecular function among all three mutants (*let-418, sin-3*, and *spr-1*), two mutants (*sin-3* and *spr-1*), and the *let-418* mutant are shown.

## References

1. Cavalli G. Chromatin and epigenetics in development: blending cellular memory with cell fate plasticity. Development (Cambridge, England). 2006;133(11):2089-94. Epub 2006/05/05. doi: 10.1242/dev.02402. PubMed PMID: 16672331.

2. Garcia-Ramirez M, Rocchini C, Ausio J. Modulation of chromatin folding by histone acetylation. The Journal of biological chemistry. 1995;270(30):17923-8. Epub 1995/07/28. PubMed PMID: 7629098.

3. Cosgrove MS, Boeke JD, Wolberger C. Regulated nucleosome mobility and the histone code. Nature structural & molecular biology. 2004;11(11):1037-43. Epub 2004/11/04. doi: 10.1038/nsmb851. PubMed PMID: 15523479.

4. Vaquero A, Sternglanz R, Reinberg D. NAD+-dependent deacetylation of H4 lysine 16 by class III HDACs. Oncogene. 2007;26(37):5505-20. Epub 2007/08/19. doi: 10.1038/sj.onc.1210617. PubMed PMID: 17694090.

5. Yang XJ, Seto E. The Rpd3/Hda1 family of lysine deacetylases: from bacteria and yeast to mice and men. Nature reviews Molecular cell biology. 2008;9(3):206-18. Epub 2008/02/23. doi: 10.1038/nrm2346. PubMed PMID: 18292778; PubMed Central PMCID: PMCPMC2667380.

6. Hayakawa T, Nakayama J. Physiological roles of class I HDAC complex and histone demethylase. Journal of biomedicine & biotechnology. 2011;2011:129383. Epub 2010/11/05. doi: 10.1155/2011/129383. PubMed PMID: 21049000; PubMed Central PMCID: PMCPMC2964911.

7. Feng D, Liu T, Sun Z, Bugge A, Mullican SE, Alenghat T, et al. A circadian rhythm orchestrated by histone deacetylase 3 controls hepatic lipid metabolism. Science (New York, NY). 2011;331(6022):1315-9. Epub 2011/03/12. doi: 10.1126/science.1198125. PubMed PMID: 21393543; PubMed Central PMCID: PMCPMC3389392.

8. Knutson SK, Chyla BJ, Amann JM, Bhaskara S, Huppert SS, Hiebert SW. Liver-specific deletion of histone deacetylase 3 disrupts metabolic transcriptional networks. The EMBO journal. 2008;27(7):1017-28. Epub 2008/03/21. doi: 10.1038/emboj.2008.51. PubMed PMID: 18354499; PubMed Central PMCID: PMCPMC2323257.

9. Miller KM, Tjeertes JV, Coates J, Legube G, Polo SE, Britton S, et al. Human HDAC1 and HDAC2 function in the DNA-damage response to promote DNA nonhomologous end-joining. Nature structural & molecular biology. 2010;17(9):1144-51. Epub 2010/08/31. doi: 10.1038/nsmb.1899. PubMed PMID: 20802485; PubMed Central PMCID: PMCPMC3018776.

10. Sun Z, Miller RA, Patel RT, Chen J, Dhir R, Wang H, et al. Hepatic Hdac3 promotes gluconeogenesis by repressing lipid synthesis and sequestration. Nature medicine. 2012;18(6):934-42. Epub 2012/05/09. doi: 10.1038/nm.2744. PubMed PMID: 22561686; PubMed Central PMCID: PMCPMC3411870.

11. Wilting RH, Yanover E, Heideman MR, Jacobs H, Horner J, van der Torre J, et al. Overlapping functions of Hdac1 and Hdac2 in cell cycle regulation and haematopoiesis. The EMBO journal. 2010;29(15):2586-97. Epub 2010/06/24. doi: 10.1038/emboj.2010.136. PubMed PMID: 20571512; PubMed Central PMCID: PMCPMC2928690.

12. Mannervik M, Levine M. The Rpd3 histone deacetylase is required for segmentation of the Drosophila embryo. Proceedings of the National Academy of Sciences of the United States of America. 1999;96(12):6797-801. Epub 1999/06/09. PubMed PMID: 10359792; PubMed Central PMCID: PMCPMC21995.

13. Montgomery RL, Davis CA, Potthoff MJ, Haberland M, Fielitz J, Qi X, et al. Histone deacetylases 1 and 2 redundantly regulate cardiac morphogenesis, growth, and contractility. Genes & development. 2007;21(14):1790-802. Epub 2007/07/20. doi: 10.1101/gad.1563807. PubMed PMID: 17639084; PubMed Central PMCID: PMCPMC1920173.

14. Shi Y, Mello C. A CBP/p300 homolog specifies multiple differentiation pathways in Caenorhabditis elegans. Genes & development. 1998;12(7):943-55. Epub 1998/05/09. doi: 10.1101/gad.12.7.943. PubMed PMID: 9531533; PubMed Central PMCID: PMCPMC316678.

15. Vecera J, Bartova E, Krejci J, Legartova S, Komurkova D, Ruda-Kucerova J, et al. HDAC1 and HDAC3 underlie dynamic H3K9 acetylation during embryonic neurogenesis and in schizophrenia-like animals. Journal of cellular physiology. 2018;233(1):530-48. Epub 2017/03/17. doi: 10.1002/jcp.25914. PubMed PMID: 28300292.

16. Genome sequence of the nematode C. elegans: a platform for investigating biology. Science (New York, NY). 1998;282(5396):2012-8. Epub 1998/12/16. PubMed PMID: 9851916.

17. Sulston JE, Schierenberg E, White JG, Thomson JN. The embryonic cell lineage of the nematode Caenorhabditis elegans. Developmental biology. 1983;100(1):64-119. Epub 1983/11/01. PubMed PMID: 6684600.

18. Wenzel D, Palladino F, Jedrusik-Bode M. Epigenetics in C. elegans: facts and challenges. Genesis (New York, NY: 2000). 2011;49(8):647-61. Epub 2011/05/04. doi: 10.1002/dvg.20762. PubMed PMID: 21538806.

19. Ranawade AV, Cumbo P, Gupta BP. Caenorhabditis elegans histone deacetylase hda-1 is required for morphogenesis of the vulva and LIN-12/Notch-mediated specification of uterine cell fates. G3 (Bethesda, Md). 2013;3(8):1363-74. Epub 2013/06/26. doi: 10.1534/g3.113.006999. PubMed PMID: 23797102; PubMed Central PMCID: PMCPMC3737176.

20. Bender AM, Kirienko NV, Olson SK, Esko JD, Fay DS. lin-35/Rb and the CoREST ortholog spr-1 coordinately regulate vulval morphogenesis and gonad development in C. elegans. Developmental biology. 2007;302(2):448-62. Epub 2006/10/31. doi: 10.1016/j.ydbio.2006.09.051. PubMed PMID: 17070797; PubMed Central PMCID: PMCPMC1933485.

21. Choy SW, Wong YM, Ho SH, Chow KL. C. elegans SIN-3 and its associated HDAC corepressor complex act as mediators of male sensory ray development. Biochemical and biophysical research communications. 2007;358(3):802-7. Epub 2007/05/18. doi: 10.1016/j.bbrc.2007.04.194. PubMed PMID: 17506990.

22. Solari F, Ahringer J. NURD-complex genes antagonise Ras-induced vulval development in Caenorhabditis elegans. Current biology: CB. 2000;10(4):223-6. Epub 2000/03/08. PubMed PMID: 10704416.

23. von Zelewsky T, Palladino F, Brunschwig K, Tobler H, Hajnal A, Muller F. The C. elegans Mi-2 chromatin-remodelling proteins function in vulval cell fate determination. Development (Cambridge, England). 2000;127(24):5277-84. Epub 2000/11/15. PubMed PMID: 11076750.

24. Brenner S. The genetics of Caenorhabditis elegans. Genetics. 1974;77(1):71-94. Epub 1974/05/01. PubMed PMID: 4366476; PubMed Central PMCID: PMCPMC1213120.

25. Zhang Y, Chen D, Smith MA, Zhang B, Pan X. Selection of reliable reference genes in Caenorhabditis elegans for analysis of nanotoxicity. PloS one. 2012;7(3):e31849. Epub 2012/03/23. doi: 10.1371/journal.pone.0031849. PubMed PMID: 22438870; PubMed Central PMCID: PMCPMC3305280 members. This does not alter the authors’ adherence to all the PLoS ONE policies on sharing data and materials.

26. Kamath RS, Martinez-Campos M, Zipperlen P, Fraser AG, Ahringer J. Effectiveness of specific RNA-mediated interference through ingested double-stranded RNA in Caenorhabditis elegans. Genome biology. 2001;2(1):Research0002. Epub 2001/02/24. doi: 10.1186/gb-2000-2-1-research0002. PubMed PMID: 11178279; PubMed Central PMCID: PMCPMC17598.

27. Bolger AM, Lohse M, Usadel B. Trimmomatic: a flexible trimmer for Illumina sequence data. Bioinformatics (Oxford, England). 2014;30(15):2114-20. Epub 2014/04/04. doi: 10.1093/bioinformatics/btu170. PubMed PMID: 24695404; PubMed Central PMCID: PMCPMC4103590.

28. Love MI, Huber W, Anders S. Moderated estimation of fold change and dispersion for RNA-seq data with DESeq2. Genome biology. 2014;15(12):550. Epub 2014/12/18. doi: 10.1186/s13059-014-0550-8. PubMed PMID: 25516281; PubMed Central PMCID: PMCPMC4302049.

29. Nomoto Y, Kubota Y, Ohnishi Y, Kasahara K, Tomita A, Oshime T, et al. Gene Cascade Finder: A tool for identification of gene cascades and its application in Caenorhabditis elegans. PloS one. 2019;14(9):e0215187. Epub 2019/09/11. doi: 10.1371/journal.pone.0215187. PubMed PMID: 31504044; PubMed Central PMCID: PMCPMC6736238.

30. Dennis G, Sherman BT, Hosack DA, Yang J, Gao W, Lane HC, et al. DAVID: Database for Annotation, Visualization, and Integrated Discovery. Genome Biology. 2003;4(9):R60. doi: 10.1186/gb-2003-4-9-r60.

31. Passannante M, Marti CO, Pfefferli C, Moroni PS, Kaeser-Pebernard S, Puoti A, et al. Different Mi-2 complexes for various developmental functions in Caenorhabditis elegans. PloS one. 2010;5(10):e13681. Epub 2010/11/10. doi: 10.1371/journal.pone.0013681. PubMed PMID: 21060680; PubMed Central PMCID: PMCPMC2965115.

32. Dufourcq P, Victor M, Gay F, Calvo D, Hodgkin J, Shi Y. Functional requirement for histone deacetylase 1 in Caenorhabditis elegans gonadogenesis. Molecular and cellular biology. 2002;22(9):3024-34. Epub 2002/04/10. PubMed PMID: 11940660; PubMed Central PMCID: PMCPMC133761.

33. Ma P, Schultz RM. HDAC1 and HDAC2 in mouse oocytes and preimplantation embryos: Specificity versus compensation. Cell death and differentiation. 2016;23(7):1119-27. Epub 2016/04/16. doi: 10.1038/cdd.2016.31. PubMed PMID: 27082454; PubMed Central PMCID: PMCPMC4946893.

34. Pillai R, Coverdale LE, Dubey G, Martin CC. Histone deacetylase 1 (HDAC-1) required for the normal formation of craniofacial cartilage and pectoral fins of the zebrafish. Developmental dynamics: an official publication of the American Association of Anatomists. 2004;231(3):647-54. Epub 2004/09/18. doi: 10.1002/dvdy.20168. PubMed PMID: 15376317.

35. Whetstine JR, Ceron J, Ladd B, Dufourcq P, Reinke V, Shi Y. Regulation of tissue-specific and extracellular matrix-related genes by a class I histone deacetylase. Molecular cell. 2005;18(4):483-90. Epub 2005/05/17. doi: 10.1016/j.molcel.2005.04.006. PubMed PMID: 15893731.

36. Kawano T, Zheng H, Merz DC, Kohara Y, Tamai KK, Nishiwaki K, et al. C. elegans mig-6 encodes papilin isoforms that affect distinct aspects of DTC migration, and interacts genetically with mig-17 and collagen IV. Development (Cambridge, England). 2009;136(9):1433-42. Epub 2009/03/20. doi: 10.1242/dev.028472. PubMed PMID: 19297413; PubMed Central PMCID: PMCPMC2674254.

37. Huang CC, Hall DH, Hedgecock EM, Kao G, Karantza V, Vogel BE, et al. Laminin alpha subunits and their role in C. elegans development. Development (Cambridge, England). 2003;130(14):3343-58. Epub 2003/06/05. doi: 10.1242/dev.00481. PubMed PMID: 12783803.

38. Johnson RP, Kang SH, Kramer JM. C. elegans dystroglycan DGN-1 functions in epithelia and neurons, but not muscle, and independently of dystrophin. Development (Cambridge, England). 2006;133(10):1911-21. Epub 2006/04/14. doi: 10.1242/dev.02363. PubMed PMID: 16611689.

39. Kao G, Huang CC, Hedgecock EM, Hall DH, Wadsworth WG. The role of the laminin beta subunit in laminin heterotrimer assembly and basement membrane function and development in C. elegans. Developmental biology. 2006;290(1):211-9. Epub 2005/12/27. doi: 10.1016/j.ydbio.2005.11.026. PubMed PMID: 16376872.

40. Vuong-Brender TTK, Suman SK, Labouesse M. The apical ECM preserves embryonic integrity and distributes mechanical stress during morphogenesis. Development (Cambridge, England). 2017;144(23):4336-49. Epub 2017/05/21. doi: 10.1242/dev.150383. PubMed PMID: 28526752; PubMed Central PMCID: PMCPMC5769628.

41. Kim S, Wadsworth WG. Positioning of longitudinal nerves in C. elegans by nidogen. Science (New York, NY). 2000;288(5463):150-4. Epub 2001/02/07. doi: 10.1126/science.288.5463.150. PubMed PMID: 10753123.

42. Grimes JA, Nielsen SJ, Battaglioli E, Miska EA, Speh JC, Berry DL, et al. The co-repressor mSin3A is a functional component of the REST-CoREST repressor complex. The Journal of biological chemistry. 2000;275(13):9461-7. Epub 2000/03/29. doi: 10.1074/jbc.275.13.9461. PubMed PMID: 10734093.

43. Zinovyeva AY, Graham SM, Cloud VJ, Forrester WC. The C. elegans histone deacetylase HDA-1 is required for cell migration and axon pathfinding. Developmental biology. 2006;289(1):229-42. Epub 2005/11/30. doi: 10.1016/j.ydbio.2005.10.033. PubMed PMID: 16313898.

